## Supplementary Figures for "Commensal Bacteria Maintain a Qa-1^b^-restricted Unconventional CD8^+^ T Population in Gut Epithelium"

#### Figure S1

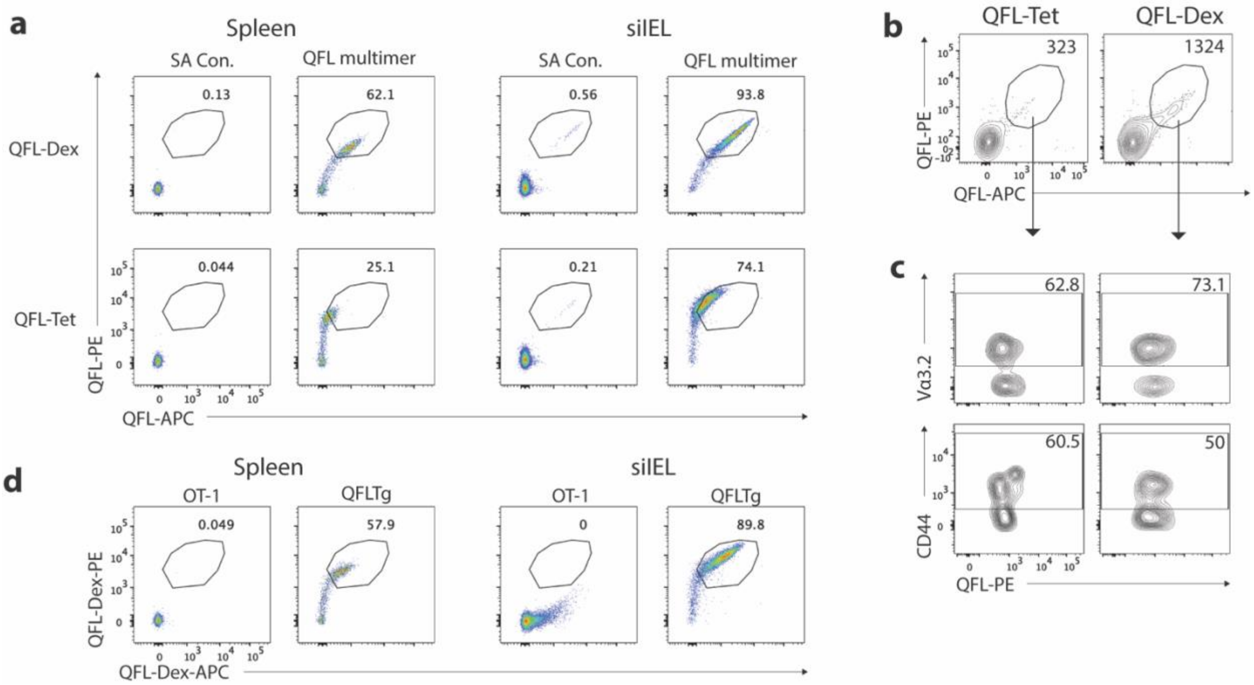


**Supplementary Fig.1 Verification of QFL-Dextramers.** (**a**) Flow cytometry of splenocytes or siIELs isolated from QFLTg mouse stained with Qa-$1^b$-FL9 tetramers (QFL-Tet) or dextramers (QFL-Dex). The unloaded streptavidin and streptavidin-dextran without QFL monomers (SA Con.) represents background signal of QFL-Tet and QFL-Dex respectively. (**b**) Flow cytometry of QFL T cells enriched from the spleen cells of naïve WT mice using QFL-Tet or QFL-Dex. Numbers in plots indicate absolute numbers of QFL-Tet/$Dex^+$ cells detected. Splenocytes from two animals were pooled and splitted into two groups for the test. (**c**) Analysis of Vα3.2 or CD44 expression on the QFL T cells enriched as in **b**. Numbers in plots indicate percentages of $V\alpha3.2^+$ or $CD44^{hi}$ cells. (**d**) Flow cytometry of splenocytes or siIELs from OT-1 or QFLTg mice stained with QFL-Dex-PE and -APC. Data are representative of 2 independent experiments.

#### Figure S2

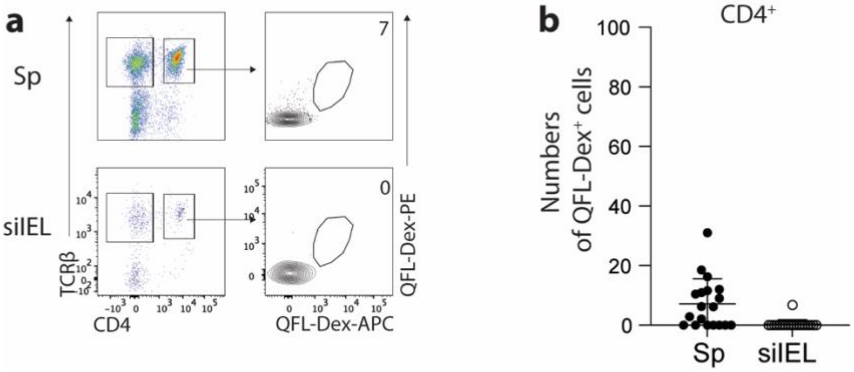


**Supplementary Fig.2 QFL T cells are barely detectable in the TCRβ$^+$CD4$^+$ population.** (**a**) Flow cytometry of QFL T cells enriched from the TCRβ$^+$CD4$^+$ population of the spleen or siIEL compartment in naïve WT mice. Numbers in plots indicate average numbers of CD4$^+$QFL-Dex$^+$ cells. (**b**) Absolute numbers of enriched CD4$^+$ QFL T cells detected as in **a**. Number of replicates is specified in the bar graphs with each symbol representing data collected from an individual mouse.

Figure S3

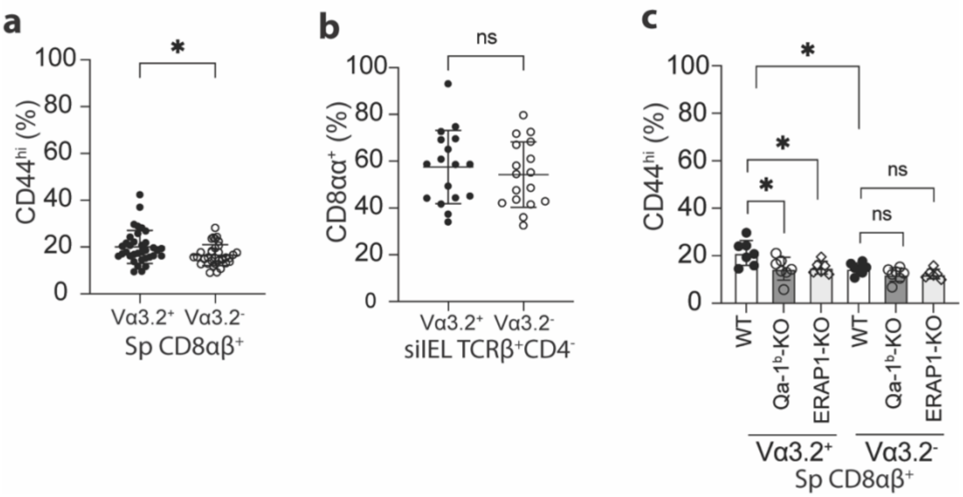


**Supplementary Fig.3 Phenotype of total Vα3.2$^+$ or Vα3.2$^-$ T population in the spleen or siIEL compartment.** (**a**) Frequencies of CD44$^{hi}$ cells detected within the total Vα3.2$^+$ or Vα3.2$^-$ CD8αβ$^+$ splenocytes from naïve WT mice**.** *P=0.0168 (**b**) Frequencies of CD8αα$^+$ cells detected within the total Vα3.2$^+$ or Vα3.2$^-$ TCRβ$^+$CD4$^-$ siIELs from naïve WT mice**.** (**c**)Frequencies of CD44$^{hi}$ cells among Vα3.2$^+$ or Vα3.2$^-$ CD8$^+$ T cells isolated from the spleen of WT, Qa-1$^b$-KO or ERAP1-KO mice. *P<0.04 Number of replicates is specified in the bar graphs with each symbol representing data collected from an individual mouse. P values were calculated with Student's *t* test. 'ns' indicates the comparison was not significant.

#### Figure S4

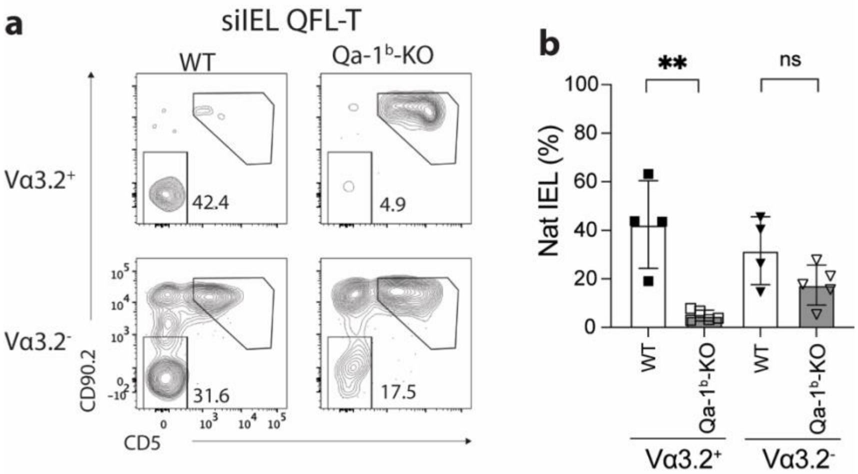


**Supplementary Fig.4 Qa-1$^{b}$-dependent expression of natIEL markers on Vα3.2$^{+}$ QFL T cells.** (**a**) Flow cytometry analysis of CD90.2 and CD5 expression on the siIEL Vα3.2$^{+}$ or Vα3.2$^{-}$ QFL-Dex$^{+}$ cells from naïve WT or Qa-1$^{b}$-KO mice. Numbers in plots indicate average percentages of CD90.2$^{-}$CD5$^{-}$ natural IEL population. (**b**) Frequencies of CD90.2$^{-}$CD5$^{-}$ cells detected as in **a**. **P=0.0023 Number of replicates is specified in the bar graphs with each symbol representing data collected from an individual mouse. P values were calculated with Student's *t* test. 'ns' indicates the comparison was not significant.

#### Figure S5

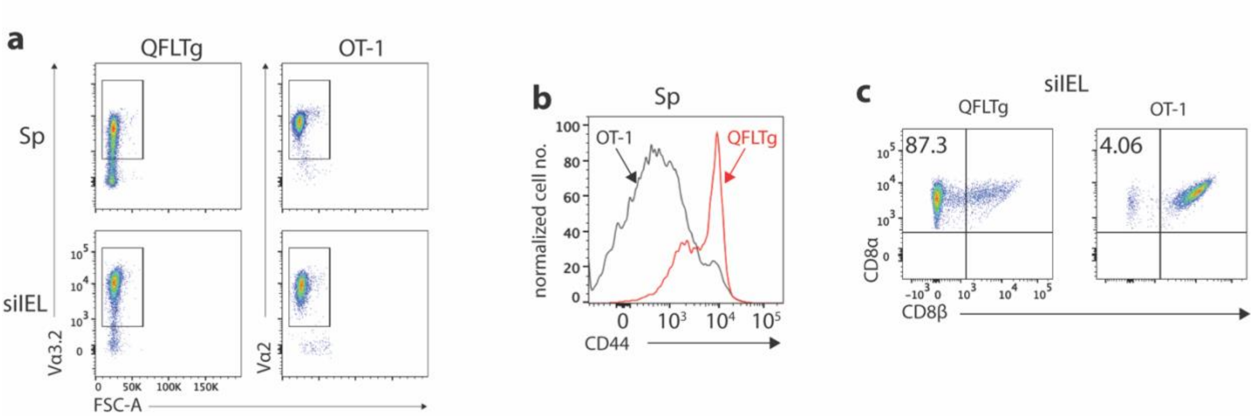


**Supplementary Fig.5 Phenotypes of QFLTg and OT-1 cells from the corresponding transgenic mice.** (**a**) Detection of QFLTg and OT-1 cells in the spleen or siIEL of QFLTg or OT-1 transgenic mice. (**b**) Flow cytometry analysis of CD44 expression on the splenic QFLTg or OT-1 cells. (**c**) Flow cytometry analysis of CD8α and CD8β expression on the siIEL QFLTg or OT-1 cells. Numbers in plots indicate percentages of CD8αα$^{+}$ cells. Results are representative of 3 independent experiments.

### Figure S6

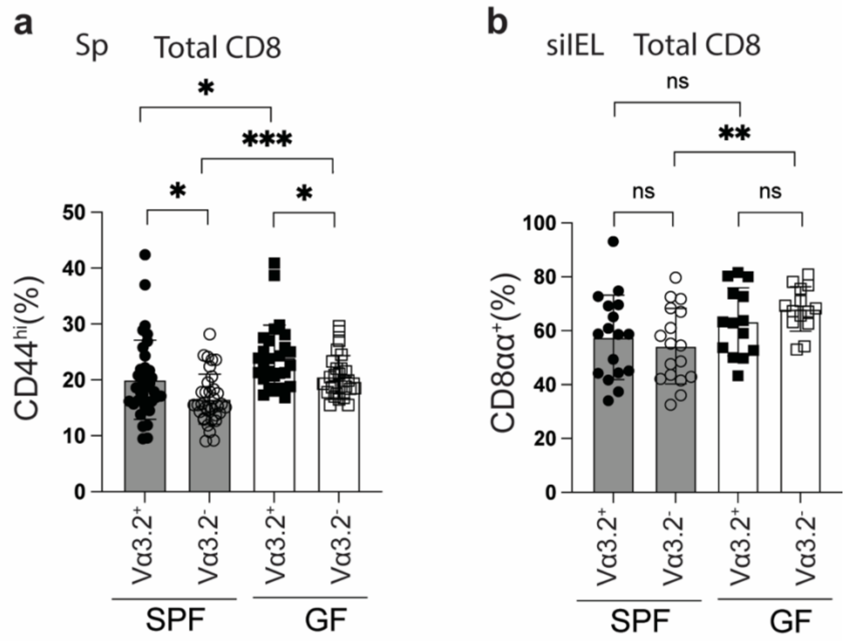


**Supplementary Fig.6 The presence of antigen experienced CD8$^+$ T cells is independent of gut microbiota.** (**a**) Frequencies of CD44$^{hi}$ cells among the total Vα3.2$^+$ or Vα3.2$^-$ TCRβ$^+$CD4$^-$ splenocytes in naïve SPF or GF WT mice. *P<0.05 ***P=0.0004 (**b**) Frequencies of CD8αα$^+$ cells among the total Vα3.2$^+$ or Vα3.2$^-$ TCRβ$^+$CD4$^-$ siIELs in naïve SPF or GF WT mice. **P=0.0030 Number of replicates is specified in the bar graphs with each symbol representing data collected from an individual mouse. P values were calculated with Student's *t* test. 'ns' indicates the comparison was not significant.

#### Figure S7

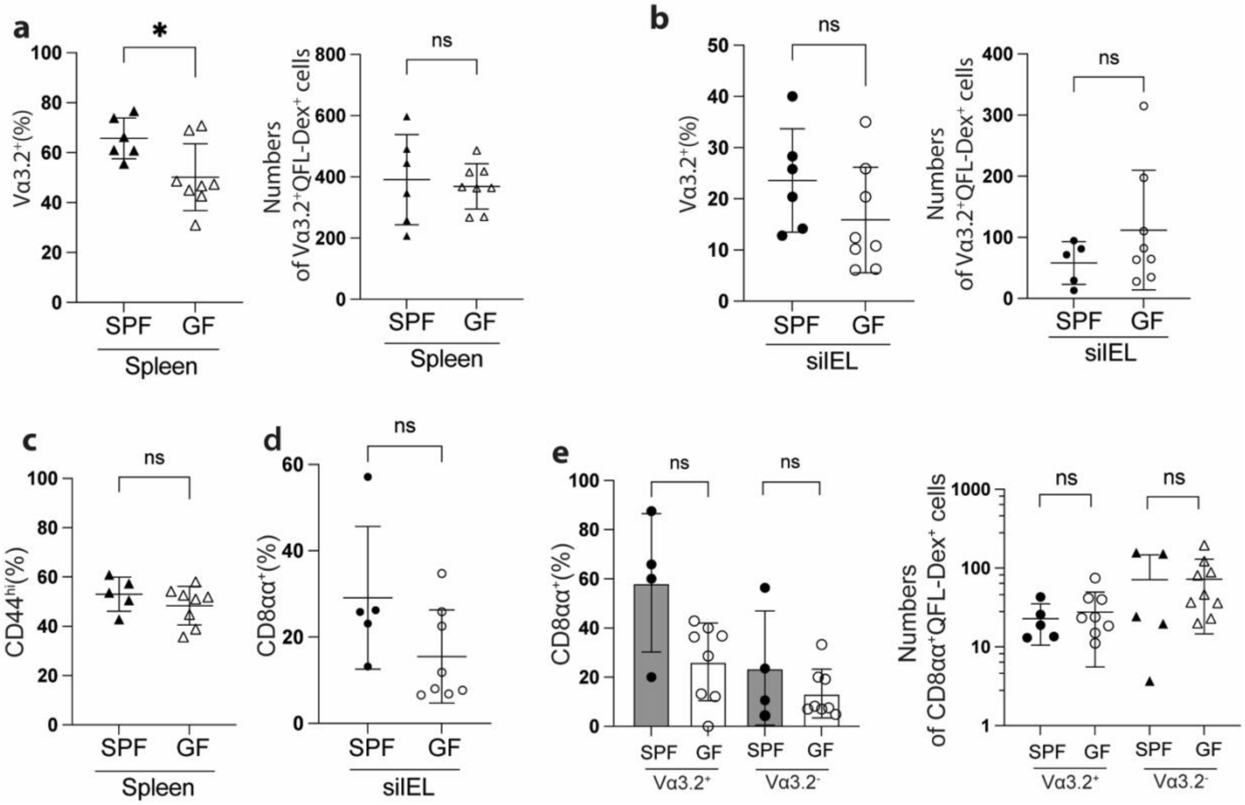


**Supplementary Fig.7 QFL T cells detected in relatively young (8~17-week) SPF or GF WT mice are phenotypically similar.** (**a**) Percentages (left) and absolute numbers (right) of Vα3.2$^+$ cells among the splenic QFL-Dex$^+$ population in SPF or GF WT mice. *P=0.0203 (**b**) Percentages (left) and absolute numbers (right) of Vα3.2$^+$ cells among siIEL QFL-Dex$^+$ population in SPF or GF WT mice. (**c**) Percentages of CD44$^{hi}$ cells among splenic QFL-Dex$^+$ population from SPF or GF WT mice. (**d**) Percentages of CD8αα$^+$ cells among siIEL QFL-Dex$^+$ population from SPF or GF WT mice. (**e**) Percentages (left) and absolute numbers (right) of CD8αα$^+$ cells among Vα3.2$^+$ or Vα3.2$^-$ QFL-Dex$^+$ population in siIEL of SPF or GF WT mice. Number of replicates is specified in the bar graphs with each symbol representing data collected from an individual mouse. P values were calculated with Student's *t* test. 'ns' indicates the comparison was not significant.

### Figure S8

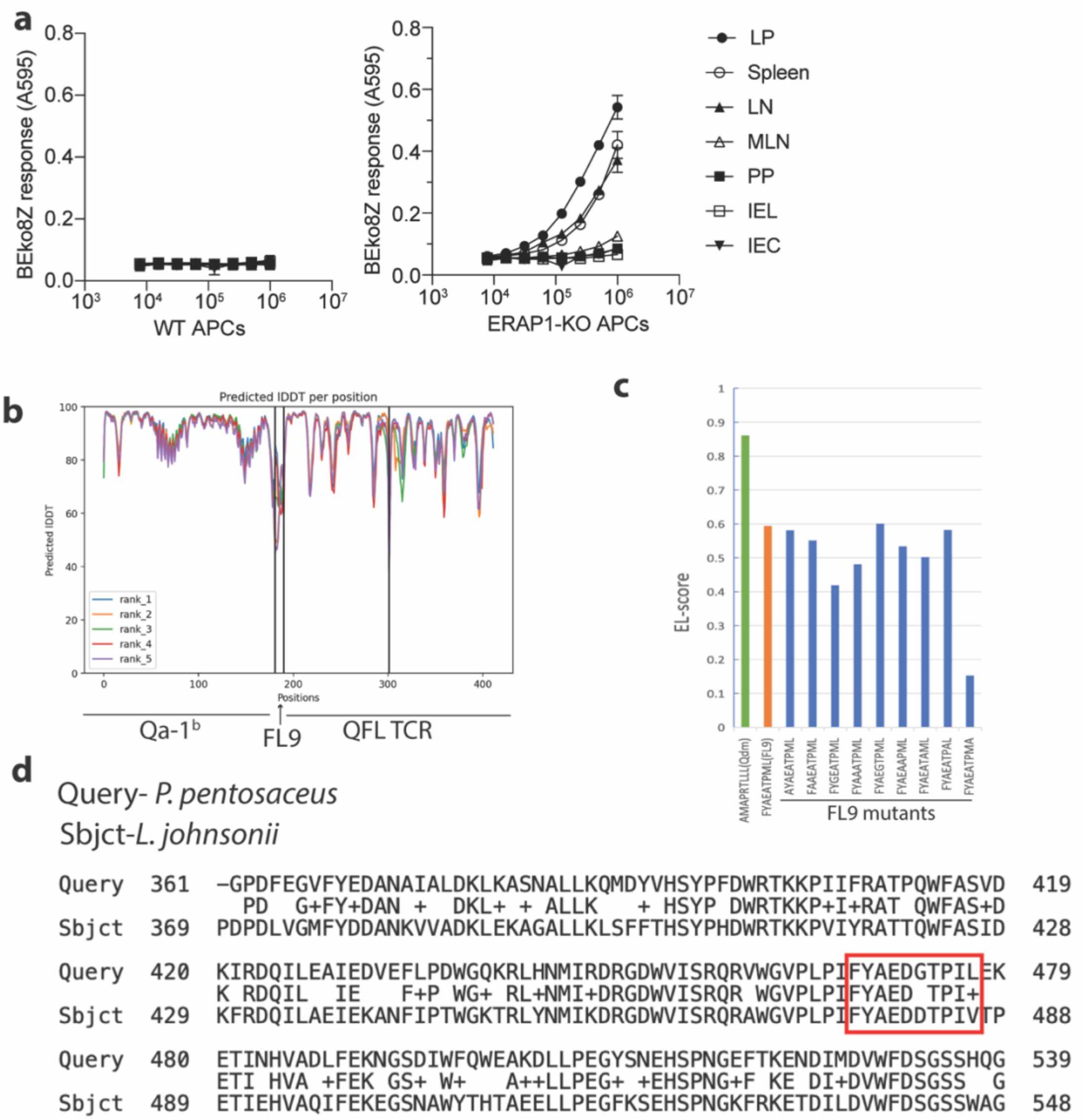


**Supplementary Fig.8 Generation of a curated library of 30 candidate FL9 homolog peptides.** (**a**) BEko8Z response to cells isolate from various tissues, including lamina propria compartment (LP), spleen, non-mesenteric lymph node (LN), mesenteric lymph node (MLN), Peyer's Patch (PP), IEL compartment or small intestinal epithelial cells (IEC) in naïve WT (left) or ERAP1-KO cells (right). (**b**) Prediction quality measured by pLDDT. The output of the highest pLDDT score ('rank_1') was selected for visualization. (**c**) Binding score of Qdm, wild-type FL9 and various FL9 mutants to Qa-1$^{b}$ predicted by NetMHCpan-4.1. (**d**) Alignment of amino acid sequences of the isoleucine tRNA ligase proteins expressed in *P. pentosaceus* or *L. johnsonii*. The FYAEDGTPIL(FL10) peptide expressed in *P. pentosaceus* and its homologue FYAEDDTPIV(FV10) expressed in *L. johnsonii* are highlighted in the red box.

Figure S9

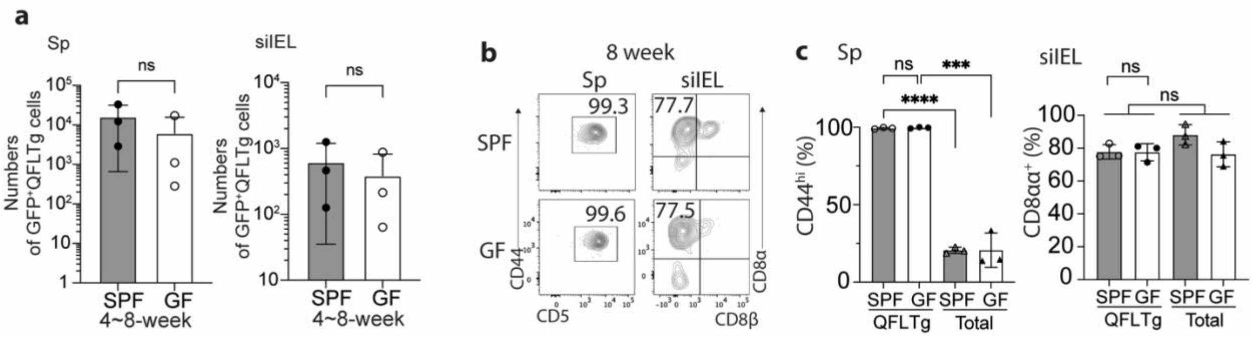


**Supplementary Fig.9 Donor-derived QFLTg cells are present and phenotypically similar in SPF or GF QFLTg_WT chimera mice before 8 weeks of age.** (**a**) Absolute numbers of donor-derived QFLTg cells detected in the spleen (left) and siIEL (right) compartment of SPF or GF QFLTg_WT chimera mice at 4~8 weeks of age. (**b**) Flow cytometry analysis of CD44 and CD5 expression on the splenic (left) or CD8α and CD8β expression on the siIEL (right) donor-derived QFLTg cells detected in 8-week SPF or GF QFLTg_WT chimera mice. Numbers in plots indicate percentages of $CD44^{hi}CD5^{+}$ or $CD8\alpha\alpha^{+}$ cells. (**c**) Frequencies of $CD44^{hi}CD5^{+}$ or $CD8\alpha\alpha^{+}$ cells among the splenic or siIEL donor-derived QFLTg populations in comparison with the total recipient-derived $TCR\beta^{+}CD4^{-}$ cells (Total) from SPF or GF QFLTg_WT chimera mice. ****P<0.0001 ***P=0.0002 Number of replicates is specified in the bar graphs with each symbol representing data collected from an individual mouse. P values were calculated with Student's *t* test. 'ns' indicates the comparison was not significant.

#### Figure S10

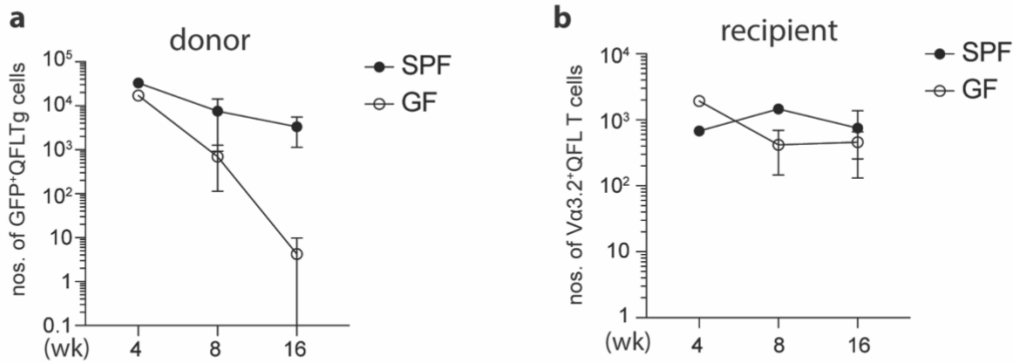


**Supplementary Fig.10 Age-associated loss of donor-derived QFLTg cells in the spleen of QFLTg_WT chimera mice.** (**a**) Absolute numbers of donor-derived $GFP^{+}$QFLTg cells enriched from the spleen of SPF or GF QFLTg_WT chimera mice at 4, 8 or 16 weeks of age. (**b**) Absolute numbers of recipient-derived $V\alpha 3.2^{+}$QFL T cells enriched from the spleen of SPF or GF QFLTg_WT chimera mice at 4, 8 or 16 weeks of age. Data is shown as mean (SD) of one 4-week SPF/GF chimera mice, two 8-week SPF/GF chimera mice and three SPF/six GF 16-week chimera mice.

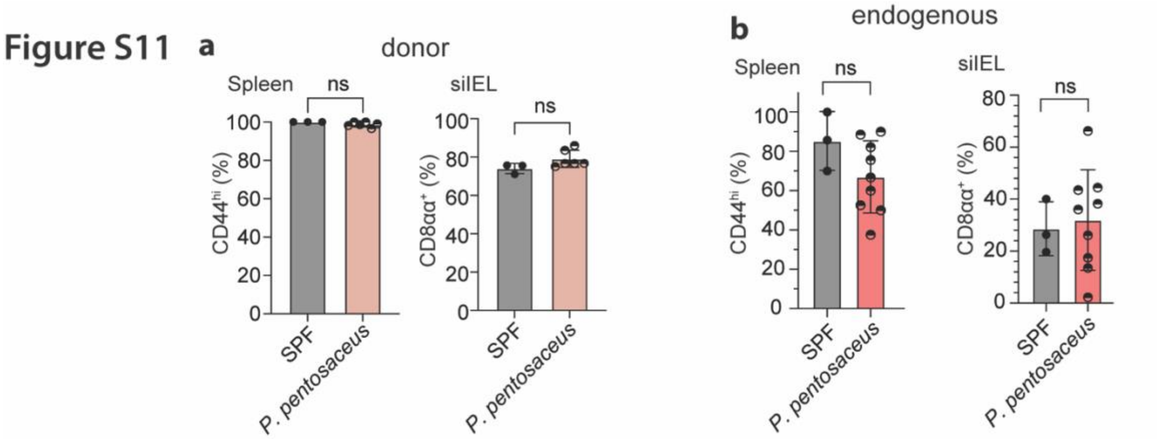


**Supplementary Fig.11 Unaltered phenotype of donor- or recipient-derived Vα3.2⁺ QFL T cells in 16-week GF chimera mice colonized with *P. pentosaceus*.** (**a**) Frequencies of $CD44^{hi}$ (left) or $CD8\alpha\alpha^{+}$(right) cells detected among the splenic or siIEL donor-derived QFLTg population from SPF QFLTg_WT chimeric mice or the GF chimeric mice colonized with *P. Pentosaceus*. (**b**) Analysis of the phenotype of endogenous $V\alpha3.2^{+}$ QFL T population as descried in **a**. Data from both QFLTg_WT chimeric mice and the non-chimeric WT mice are pooled for analysis of the endogenous populations. The number of replicates is specified in the bar graphs with each symbol representing data collected from the indicated tissue isolated from an individual mouse. P values were calculated with Student's *t* test. 'ns' indicates the comparison was not significant.

| Screening ID | NCBI ID | Peptide | Length |
|---|---|---|---|
| 1 | Q5WLW2.1_8 | YYAEAKPM | 8 |
| 2 | Q5WLW2.1_9 | YYAEAKPMV | 9 |
| 3 | A1U2V8.1_8 | FAEETPML | 8 |
| 4 | A1U2V8.1_9 | KFAEETPML | 9 |
| 5 | A6U852.1_8 | AESATPML | 8 |
| 6 | A6U852.1_9 | GAESATPML | 9 |
| 7 | A5CDU2.1_8 | YHQATPML | 8 |
| 8 | A5CDU2.1_9 | QYHQATPML | 9 |
| 9 | Q2G5G2.1 | REAEATAML | 9 |
| 10 | A4SR98.1 | FYAEEHATP | 9 |
| 11 | Q03EY9.1 | FYAEDGTPIL | 10 |
| 12 | A4W6X4.1 | FHAEATPL | 8 |
| 13 | A3M02.2 | FFAEAKRML | 9 |
| 14 | Q15W53.1_8 | FYEACPML | 8 |
| 15 | Q15W53.1_9 | SFYEACPML | 9 |
| 16 | P71338.2 | YAAATPYALML | 11 |
| 17 | A9KDQ8.1 | YAEATSGLPLL | 11 |
| 18 | Q12C38.1 | NAAEATRML | 9 |
| 19 | P55386.1 | FVIEASPML | 9 |
| 20 | B8H8S3.1 | GYADAKPMV | 9 |
| 21 | P66878.1 | FYADATERI | 9 |
| 22 | A1WN86.1 | FYADVTPAE | 9 |
| 23 | Q1QF31.1 | FYTAELTPLML | 11 |
| 24 | A9BNI6.1 | IEAIATPML | 9 |
| 25 | A0LJA8.1 | FYAGATGMM | 9 |
| 26 | B8F5T7.1 | AVTEASPML | 9 |
| 27 | A9WSC4.1 | FYAEAGGQV | 9 |
| 28 | A7IFD4.1 | YYDEATAMV | 9 |
| 29 | Q1RGF1.1_10 | FYGDATRMDL | 10 |
| 30 | Q1RGF1.1_9 | FYGDATRMD | 9 |

**Supplementary Table.1 Library of the 30 candidate FL9 homolog peptides expressed in commensal bacteria.**
